# Expression analysis of colon adenocarcinoma in the South Indian population reveals novel signals of pathogenesis

**DOI:** 10.1101/2020.10.14.338814

**Authors:** Prasanth S. Ariyannur, Reenu Anne Joy, Veena Menon, Roopa Rachael Paulose, Keechilat Pavithran, Damodaran M. Vasudevan

## Abstract

High throughput somatic expression analyses of colon adenocarcinoma conducted so far were mostly in the western population, and no major studies are currently available in the Indian population. We have performed Nanostring PanCancer pathway panel assay in Stage II colon cancer (n = 11) and compared against normal colon tissues (n = 4). Differentially expressed (DE) genes were identified and superimposed on the Cancer Genome Atlas (TCGA) data, from the Genome Expression Profiling Interactive Analysis (GEPIA) and Tumor Immune Estimation Resource (TIMER).83 out of 730 genes were significantly different (*p*-value < 0.01), 19 of which had a fold-change |FC(log_2_)| ≥ 2. A comparison of these signals on TCGA COAD data revealed four common (MET, MCM2, ETV4, and MMP7) and15 uncommon genes. On group comparison, ETV4 expression was significantly higher in microsatellite stable (MSS). Significant DE genes, unique in the study, were INHBA, COL1A1, COL11A1, COMP, SFRP4, and SPP1, which were clustered in STRING.db network analysis and correlated with tumor-infiltrating immune cells in TIMER.MMR functions of colon cancer pathogenesis in India may be differentiated by validating the somatic expression of most common genes identified in the study.

## Introduction

More than thirty years of research on molecular pathophysiology of colorectal cancer (CRC) has led to the identification of many novel mechanisms of tumorigenesis. Based on whole-genome sequencing studies, colon adenocarcinoma (COAD) has the highest number of overall general genomic alterations per tumor compared to any other tumor tissue types, irrespective of the stage of the disease^1^. This was explained by the rapid self-renewing nature of normal colonic epithelial cells, the overall additive effect of a series of small driver mutations among the background of passenger mutations creating a survival advantage. In the western population, studies have shown that overall pathogenicity of CRC is primarily due to chromosomal instability (CIN) leading to widespread loss of heterozygosity (LOH) and gross chromosomal rearrangements and high somatic copy number variations ^2,3^. Large cohort studies done in the western population attributed the occurrence of CRC primarily to sporadic multi-molecular genetic changes (70-80%), causing chromosome-wide changes ^4^, which led to the classification of CRC into distinct molecular subtypes^5,6^.Defective DNA mismatch repair (MMR) and consequent microsatellite instability (MSI), classified as hypermutated type in The Cancer Genome Atlas (TGCA),are the second common cause, accounting for roughly one-fifth of sporadic molecular genetic changes^5^. Less than 5% of CRC was found to be due to delineable hereditary molecular factors, mainly autosomal dominant inherited multi-organ cancer disorder, Lynch Syndrome (LS), due to constitutional and inherited mutations in four MMR genes-*MLH1, MSH2, MSH6, PMS2*,and*EPCAM*^7^. Deficient MMR comprises approximately 15% of the CRC pathogenesis, including both sporadic and hereditary causes in the western population ^8^.

The complexity of MMR function due to the involvement of more than 30 genes ^9^, epigenetic inactivation ^10^, microRNA mediated regulation of MMR gene transcripts ^11^, and hetero-oligomeric co-dependence of MMR proteins ^12^, leading to variable phenotypic penetrance are some of the very few known factors for diagnostic and scientific challenges of this syndrome. The exorbitant diagnostic cost and subsequent lack of compliance for repeated investigations are explained partially by these challenges. However, multiple levels of diagnostic maneuvers to circumvent these issues led to a complex, expensive, indirect, and often inadequate clinical conclusion. All of these concerns suggest the lack of a better diagnostic algorithm and perhaps better biomarker(s) suited for different population create a great deal of diagnostic lacuna in the local population.

The prevalence rate of colon cancer in India, according to a recent population-based cancer registry by the Indian Council for Medical Research (ICMR), is on the rise. Average prevalence rates of 4.3 and 3.4 in 100,000 males and females respectively, across different parts of India, are further increasing, as per the recent epidemiological reports ^13^. The International Association of Cancer Registries (IACR) Global Cancer Observatory (GLOBOCAN) showed an incidence of 5.8 − 9 per 100,000, with a slightly higher rate of mortality in men than women (4.1 − 5.8 vs 3.5 – 5.3).Large studies on DNA microsatellites in different cancer types, by TCGA, did not include representative people from major Indian ethnic groups ^14^. Pioneer studies, conducted by a center in Tamil Nadu, on the Indian CRC showed that 67% of cases had deficient MMR^15^. Another study done in Karnataka showed a prevalence of about 10% ^16^. Irrespective of the pathways involved in the pathogenesis of early or late-onset CRC, MSI was found to be around 40% in another study conducted in Andhra Pradesh ^17^. More recent studies from the same region showed 48% of cases were MSI ^18^. There have been not many studies conducted in India to address the reason behind the high prevalence of MMR deficiency occurring in India compared to other populations. Furthermore, by immunohistochemical study conducted in the our population, during the last eight years, identified deficient MMR in about 27% cases of Stage II CRC ^9^. However, there has not been any molecular pathogenesis study to explore the reasons behind higher prevalence of MMR in CRC pathogenesis in Indian population.

To address this, we selected a set of Stage II CRC from a tertiary cancer care center, including tumors from both microsatellite stable (MSS) and unstable categories, and compared the histopathological data against a high throughput expression analysis using Nanostring nCounter Pan-Cancer pathway analysis. The expression pattern and the signals obtained from this cohort were compared against the TCGA COAD data using multiple data tools such as Genome Expression Profiling Interactive Analysis (GEPIA) and Tumor Immune Estimation Resource (TIMER)to explore the unique expression signals in CRC pathogenesis in our population.

## Results

### Samples and screening

Eleven samples of Stage II colon adenocarcinoma were selected. Tumor samples were obtained from subjects with an age of onset of 38 through 76 years (See Table 1for more details). Eight samples were from the proximal colon (ileocecal and ascending colon), two from transverse colon and one from the sigmoid colon. The tumor was previously examined by histopathology for their tumor type, differentiation, the extent of tumor invasion, lymphocyte, and associated immune cell infiltration, lympho-vascular, and perineural invasion. DNA MMR was assessed by immunohistochemical reactivity of proteins MLH1, MSH2, MSH6, and PMS2. Additionally, these samples were confirmed by MSI-PCR using two mononucleotide repeat markers (BAT25, BAT26), and three quasi-monomorphic repeat markers (NR-21, NR-24, NR-27), all of which have a high correlation with low immunoreactivity to any/all four MMR protein ^19^. Six tumor tissues had deficient MMR (both IHC and MSI −PCR), and five had proficient MMR. Five proximal colon tumors were of the MSI category. All the tumors were T3N0Mx according to the Union for International Cancer Control (UICC) staging. Six samples had moderate lymphocyte infiltration, all originating from the proximal colon region. One proximal colon tumor had severe infiltration, while three proximal and one distal had mild or no infiltration.

**Table 1:**
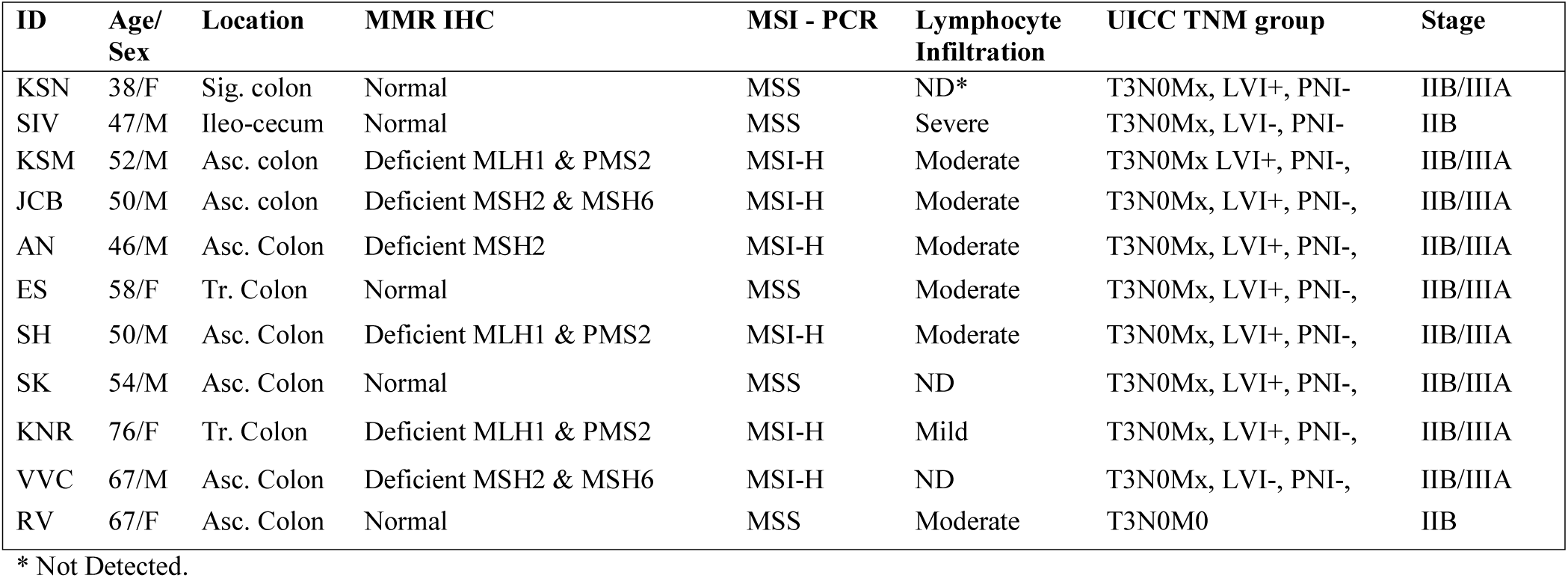
Patient and sample details

### Nanostring Expression Analysis

The sample annotation and gene expression data are provided in Supplementary Dataset File S1 and S2. Out of the 730 target signals, 83 were found to be significant (*p*-value < 0.01; FDR adj. *p*-value < 0.5) across the samples (Figure 1a). Of the 83 genes, 17 genes were upregulated, (log_2_fold-change – (FC(log_2_)) > 2) and two genes were downregulated (FC(log_2_) ≤ −2). These genes are listed in Supplementary Dataset File S3 with their corresponding *p*-values and FC(log_2_) values. On the principal component analysis (PCA), 20 genes showed segregation between tumor and control samples (Figure 1b). FC of the top 20 genes in PCA is depicted in a heatmap in Figure 1c. The 20 top genes were clustered evenly among tumor samples, and the expression pattern was reversed in the normal samples. With the Nanostring pathway analysis, signals associated with cell cycle and apoptosis, chromatin modification, and DNA damage repair were shown to be upregulated in tumor samples and downregulated in normal tissues, with few exceptions in certain samples (Figure 1d). The overall pathway score for the chromatin modification set and DNA damage repair in different samples did not correspond to their MMR status. This may be because the overall pathway effect of the whole tissue might be influenced by the tumor cells as well as Tumor-Infiltrating Lymphocytes (TILs) and macrophages, which is further understood by the GEPIA and TIMER correlation analyses.

**Figure 1:**
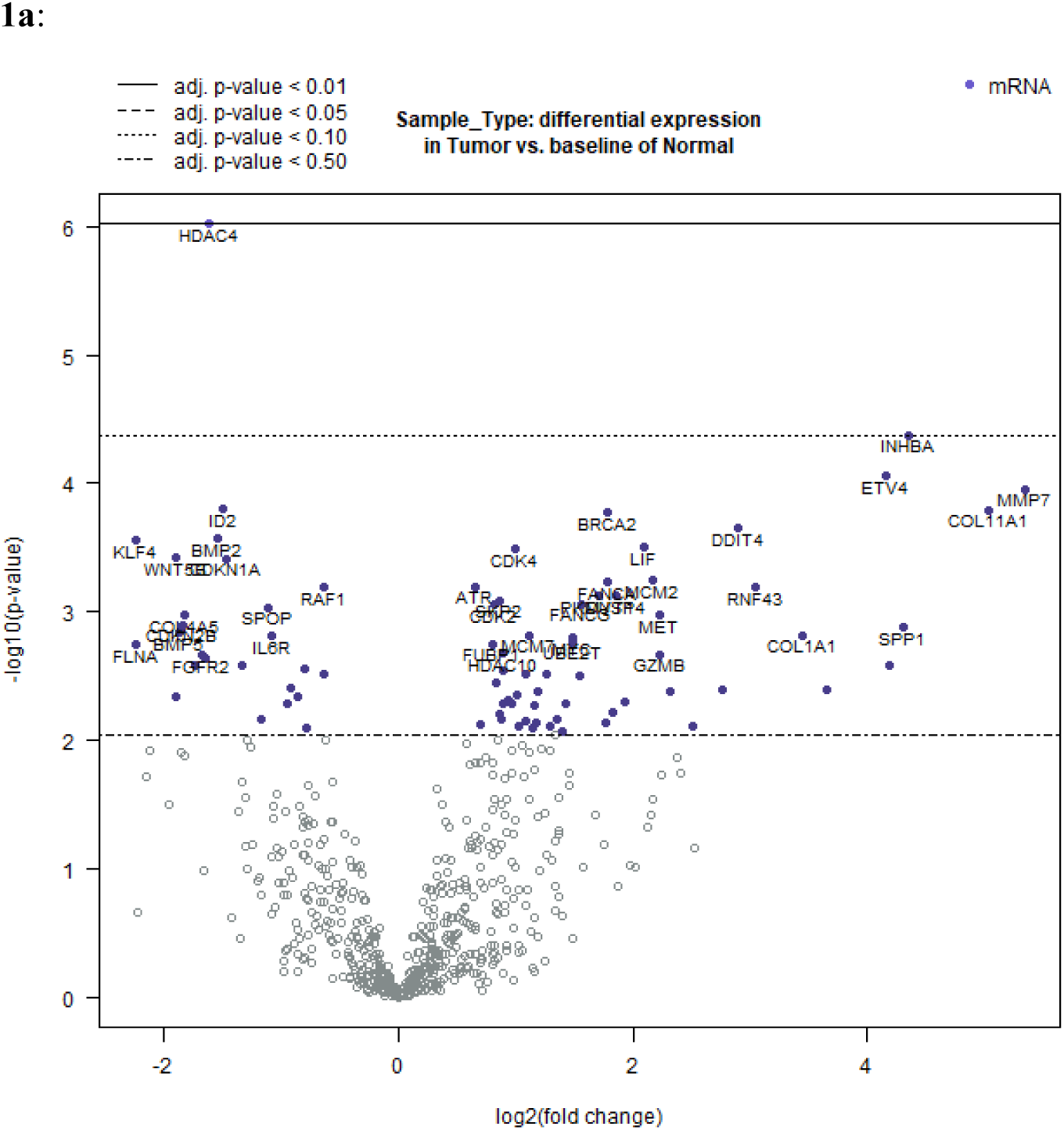
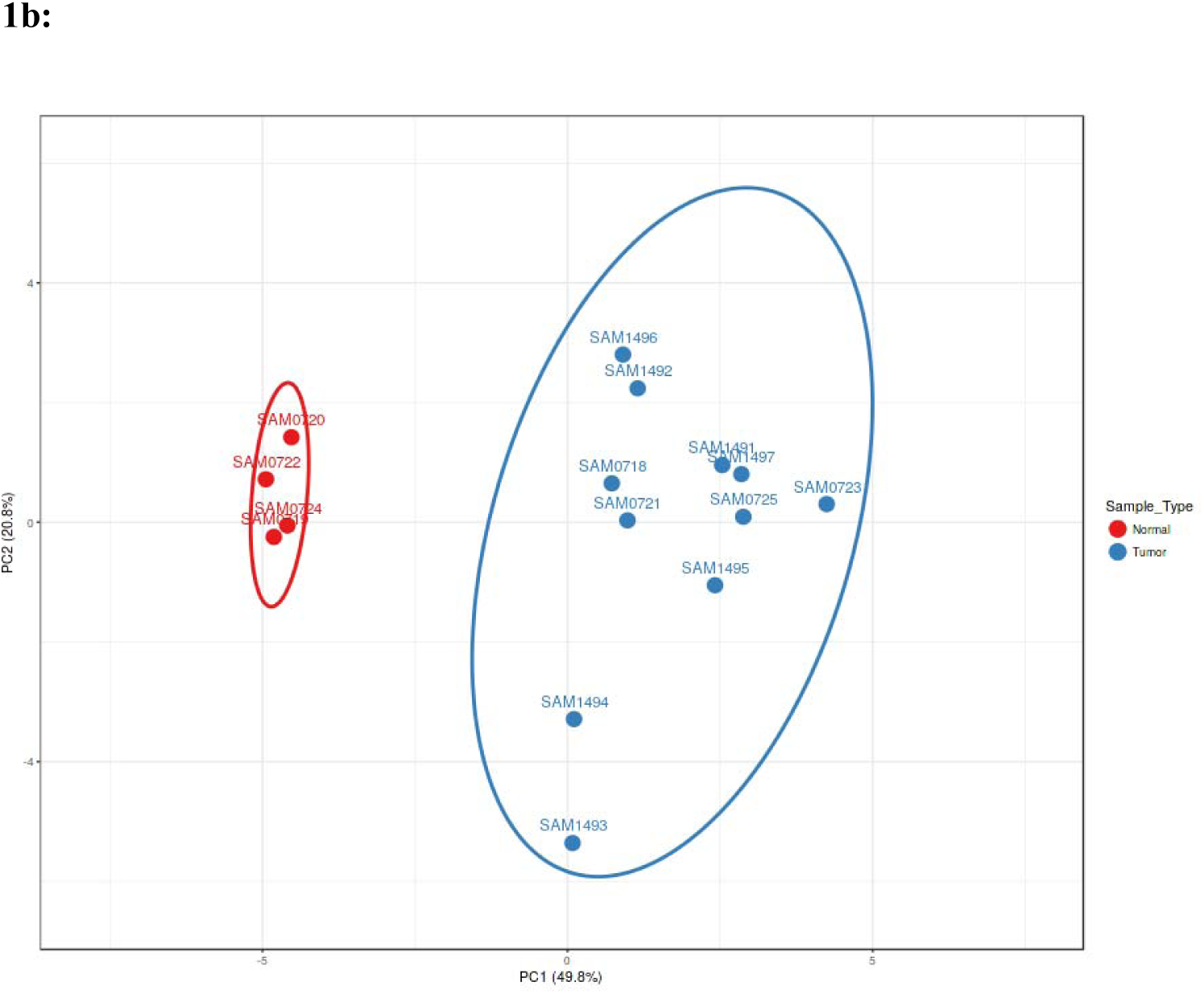
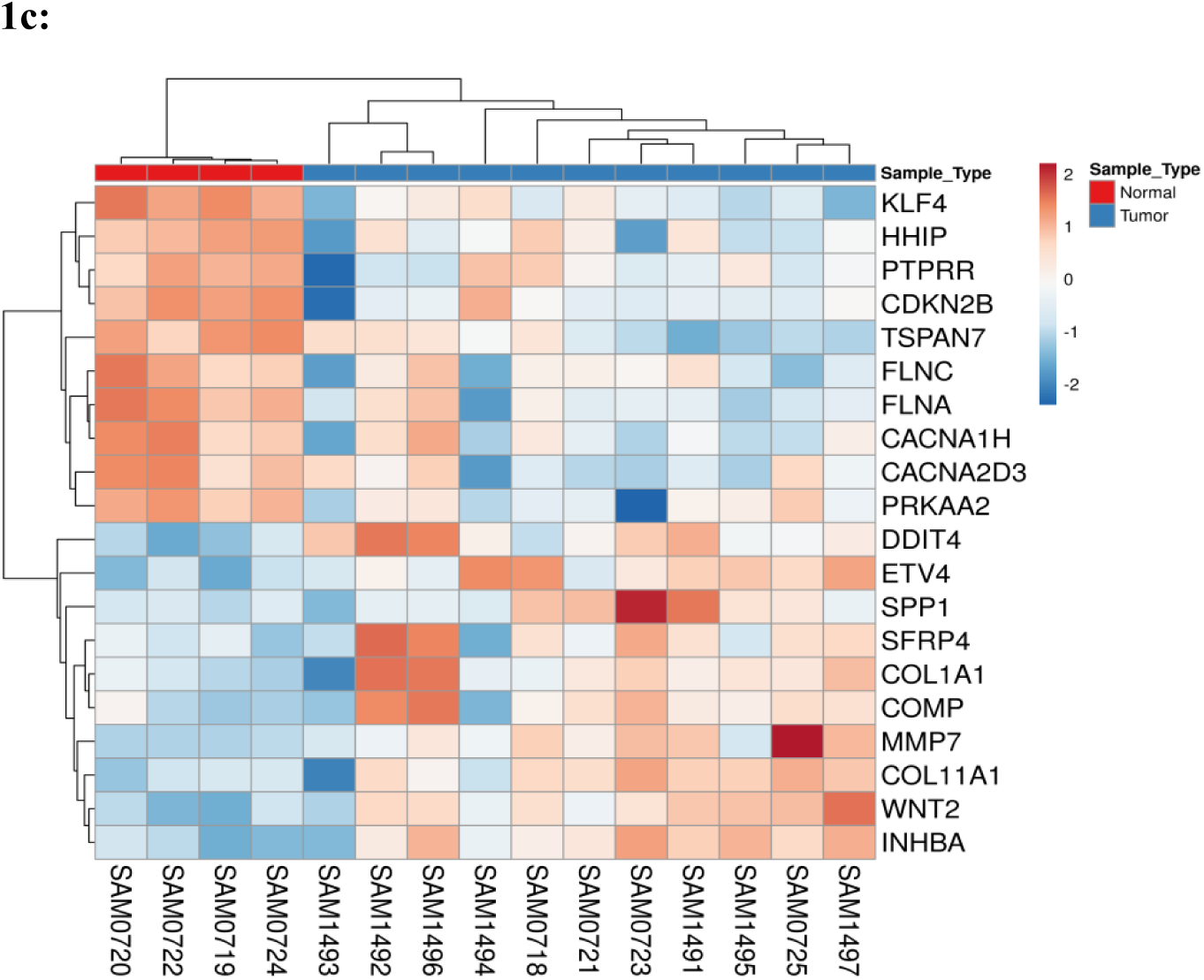
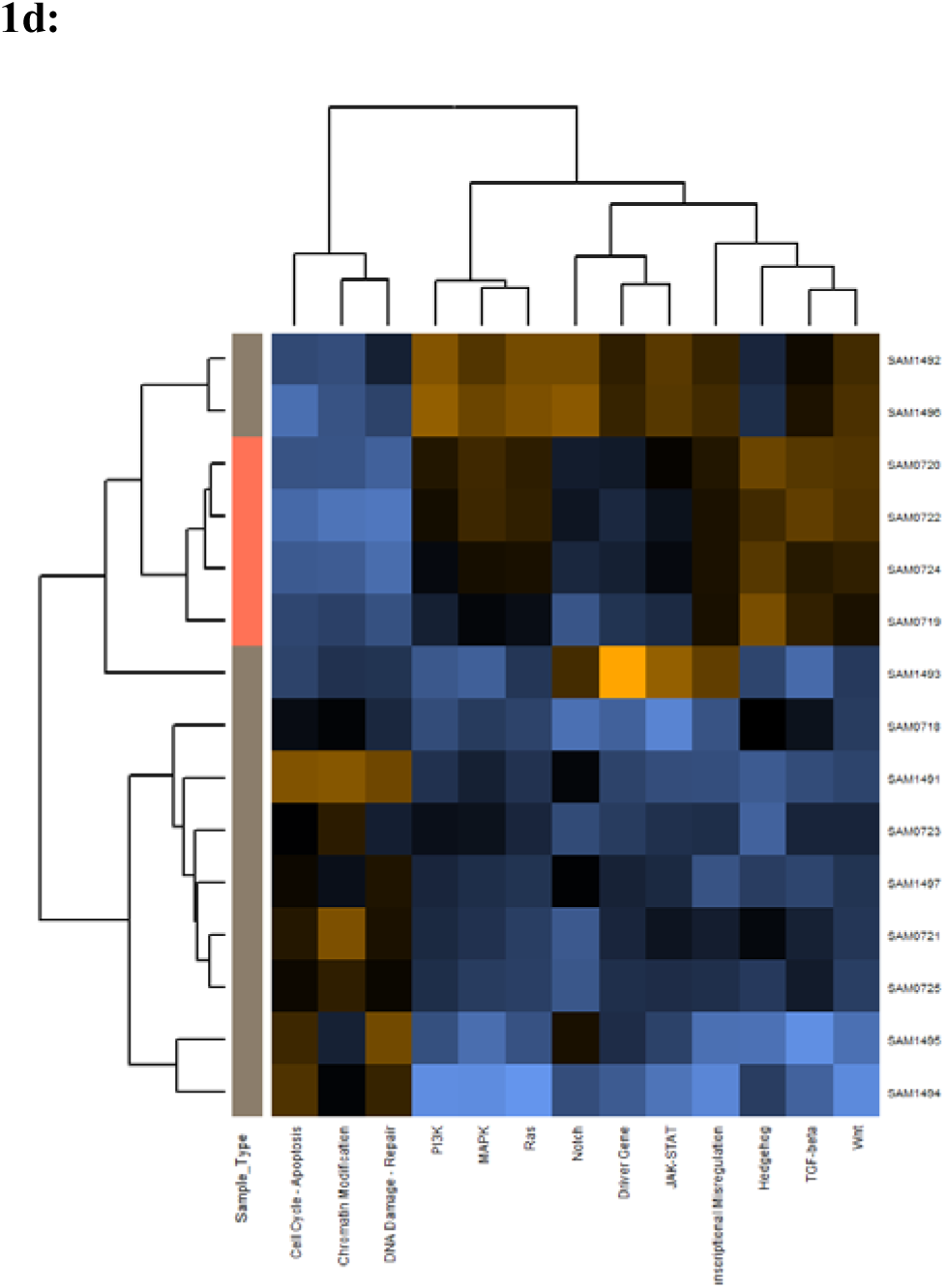
Differential expression analysis: **1a:** Volcano plot displaying the expression pattern of the entire gene set. Y-axis is −log_10_(*p*-value) and X-axis is log_2_(fold change)of each of the covariates. Significant genes fall at the top of the plot above the horizontal lines, and highly differentially expressed genes fall to either side. Horizontal lines indicate various False Discovery Rate (FDR) thresholds or p-value thresholds if there is no adjustment to the p-values. Genes are colored if the resulting *p*-value is below the given FDR or *p*-value threshold. Genes that had *p-*value < 0.01 (FDR adj. *p*-value <0.5) are 83 and among them, the 40 most statistically significant genes are labeled in the plot. **1b**: Venn diagram showing the PCA of the top 20 genes of the samples in the given cohort from the ratio data estimation. **1c**: Heatmap showing the top 20 upregulated or down-regulated genes from the differential expression analysis of the Nanostring data. All these signals have a *p*-value of < 0.01 and a log_2_fold change of at least 1. The normal and tumor samples are colored coded in the top margin. **1d**: Heatmap showing pathway score depicting an overview of how the pattern of pathway scores change across samples to understand how pathway scores cluster together and which samples exhibit similar pathway score profiles. Orange indicates high scores; blue indicates low scores. Scores are displayed on the same scale via a *Z*-transformation.

### Group comparison

To compare the effects of microsatellite instability (MSI)status and tumor immune cell infiltration (TIL) on the expression fold of mRNA signals, the expression fold change of all signals were compared separately against MSI status (MSS vs. MSI groups) and TIL status (detected or not in FFPE sections) using the Student t-test. The results are shown in Figures2&3. The expression FC of 730 genes from the Nanostring DE analysis was compared against MSI and MSS groups to reveal 13 genes that were significantly different, *p*-value < 0.05. Among these, four genes had a mean |FC(log_2_)| > 1 between the two groups (Figure 2a). Of the four, highest differential expression was shown by ETV4 (mean FC(log_2_) = 12.38; p-value = 0.035) in MSS group compared to MSI (Figure 2b). Other significant genes with FC > 1 were PLCB4 (mean FC(log_2_) = 2.43; *p*-value = 0.014), PROM1 (mean FC(log_2_) = 1.39; *p*-value = 0.025) and BIRC3 (mean FC(log_2_) = −1.13; *p*-value = 0.049). All these genes were significantly different from similar FC in expression in the TCGA as well (as shown in GEPIA analysis below). When signals were compared against groups with the two groups of TIL, the only signal that was most significant with the highest mean FC was ETV4 (mean FC(log_2_) = 12.82; p-value = 0.035) (Figure 3a&3b). Other signals that were significant between the groups, but FC(log_2_) ≈ 1were BCL2A1 (mean FC(log_2_) = −1.061, *p*-value = 0.018), CDC7 (mean FC(log_2_) = −1.42, *p*-value = 0.016), TNFRS10A (mean FC(log_2_) = −1.14, *p*-value = 0.013). Notably, ETV4 expression fold is significantly higher in TIL ND group. In both these group comparisons, ETV4 was found to be the only signal that is conspicuously higher in the MSS group with no TIL. FDR adjustment resulted in no significant difference in expression in any gene signals across either of these groups. There was no significant difference in the gene expression pattern when compared against age, sex, or anatomical location of the tumor.

**Figure 2:**
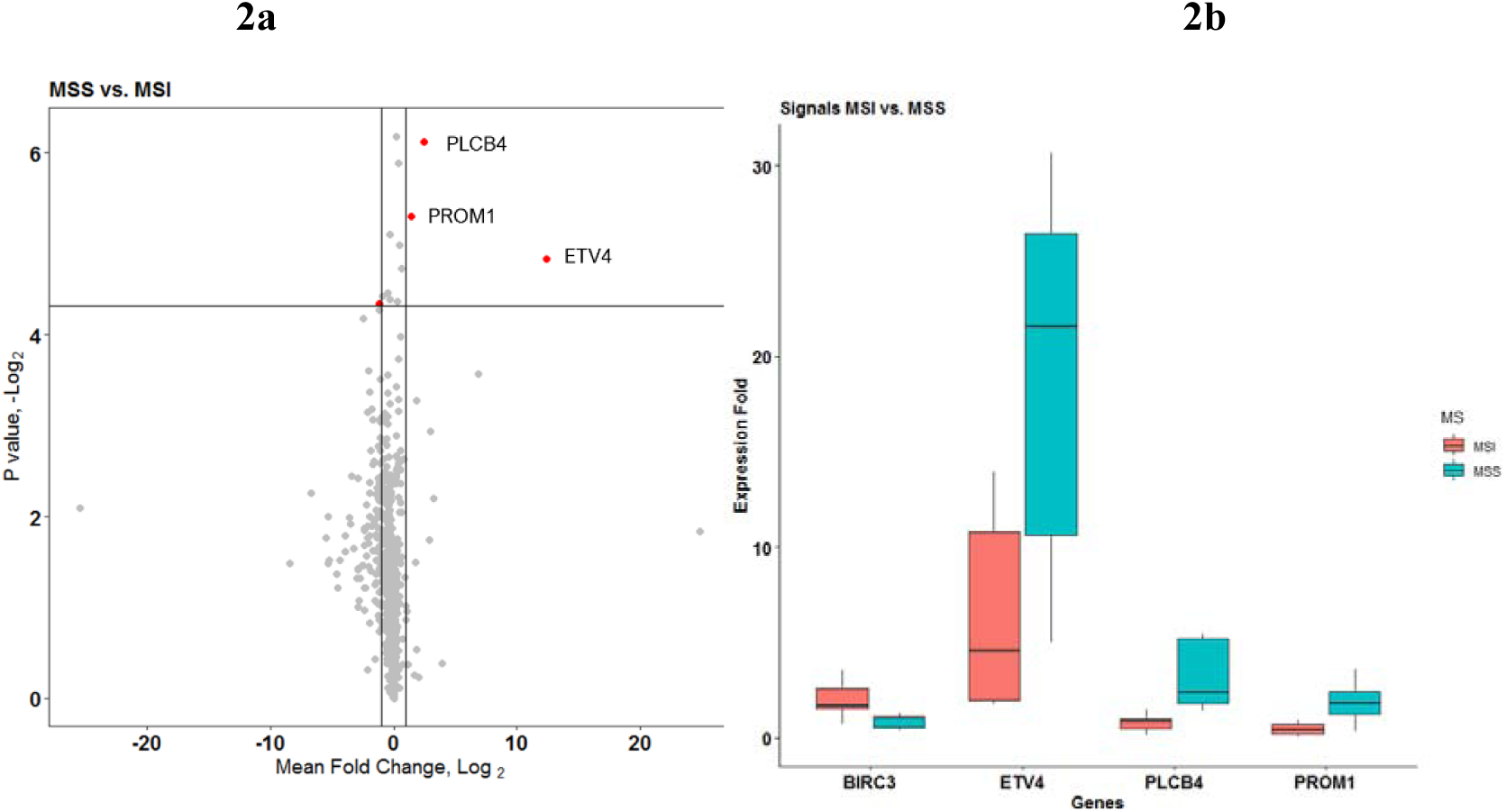
Comparison of microsatellite instability status in gene expression patterns. **2a:** Group comparison of expression fold change in MSI and MSS group using t-test. The X-axis shows the mean difference between the two groups for each of the signals and the y-axis shows the −log_2_(*p*-value). Horizontal intercept above 4 in the Y-axis is the line of significance, signals above which have a *p*-value < 0.05. The vertical line with X-intercept is at a log_2_(FC) of one. There were only three signals with |log_2_(FC)| >1 and *p*-value < 0.05. Four genes satisfied *p*-value < 0.05 and FC(log_2_) > 1-ETV4, PLCB4, and PROM1 (marked by red dot). **2b**: This figure shows the expression fold change in highly significant genes identified in figure 2a. Among them, ETV4 has the highest expression in the MSS group compared to MSI.

**Figure 3:**
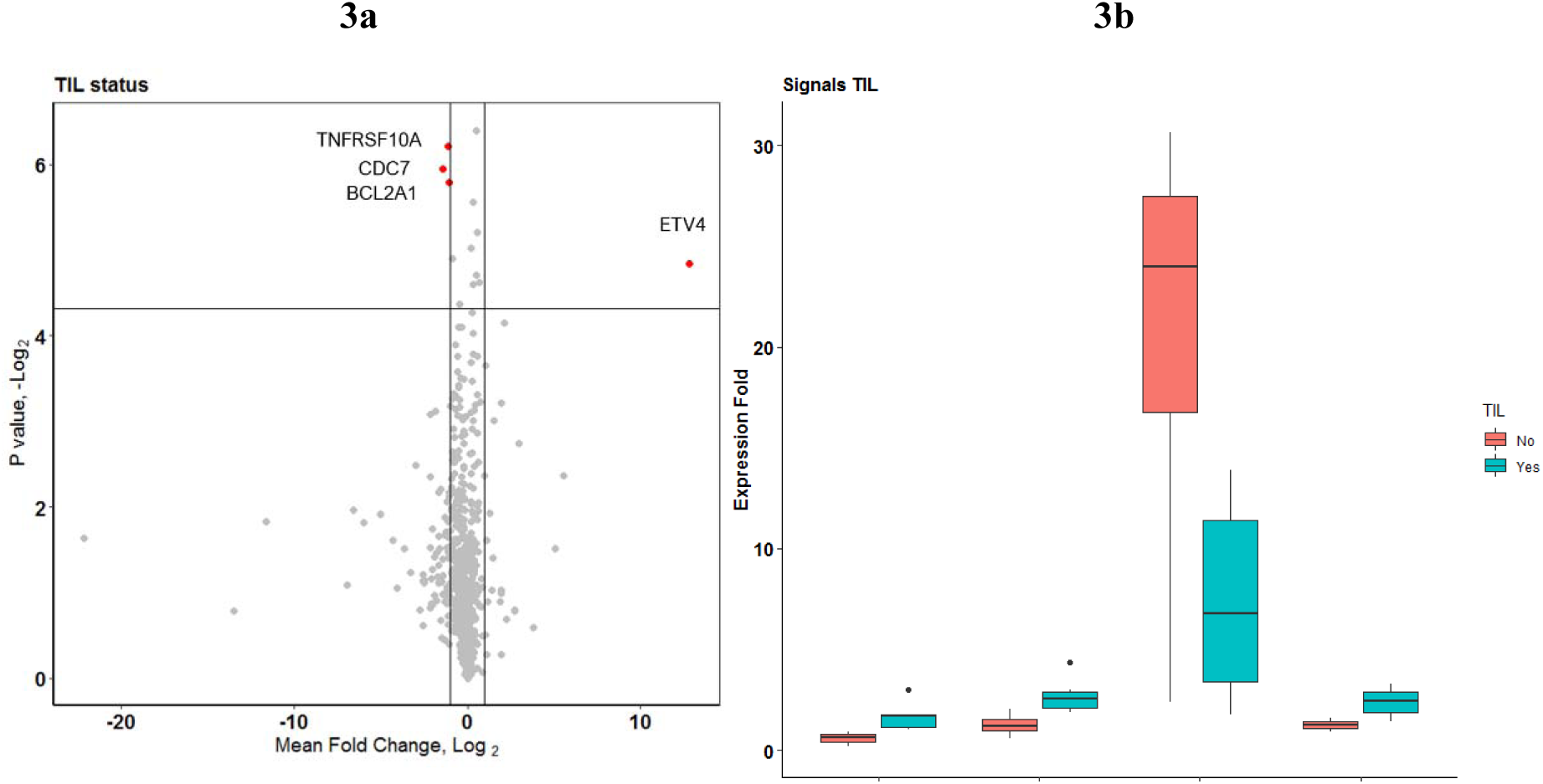
Comparison of tumor immune cell infiltration status in gene expression pattern. **3a:** Group comparison of expression fold change in TIL vs. none using t-test. Axis annotations and intercepts are the same as that of figure 2a. Four signals with |log_2_(FC)| >1 and p-value < 0.05 were ETV4, TNFRSF10A, CDC7, and BCL2A1 (marked by red dots). ETV4 is the common signal in both these categories with maximum fold change. **3b**: Boxplot depicting the expression fold change in the four genes identified in 4a. Among them, ETV4 is associated with high expression in the TIL ND group.

### TCGA COAD Correlation Analysis

#### GEPIA

The expression profile in the current study (AIMS) was compared against the TCGA COAD dataset on GEPIA. 45 out of 83 significant genes in the AIMS study were correlated with the TCGA COAD dataset (common set) in the GEPIA database (See Figure 4 and Supplementary Dataset File S3). However, 38 genes were found to be not significantly different in the TCGA dataset (unique set). The most significant gene in the TCGA from the current dataset is ETV4, while overall in the AIMS study, HDAC4 was highly significant (see volcano plot 1b). Analysis of the top 19 of the 83 signals with |FC(log_2_)| >2 revealed 14 genes were significant and five genes were from the unique set. Furthermore, eight genes with |FC(log_2_)| > 2, were significant (*p*-value < 0.01) in both AIMS study groups as well as TCGA group (common set). These genes were MMP7, COL11A1, INHBA, ETV4, RNF43, MET, GZMB, and FLNA. Among them, MMP7, ETV4,and RNF43 were found to have |FC(log_2_)| >5 in the TCGA cohort (See Supplementary Dataset File S3). Several genes that were significant in the TCGA were found to be either not significant or had FC(log_2_) < 1, in the current study. These were CCNB1, CDC6, PKMYT1, UBE2T, AINX2, ITGA2, RFC3, FANCA, MYC, IL6, and FGFR2. Signals that showed similar FC expression in both these studies were PLAU, EZH2, TNFRSF10A, BRCA2, DUSP4, MCM7, PRKDC, and RIN1. One signal, HRAS, showed an upregulation, FC(log_2_)= 1.01 in our study and down-regulation in TCGA GEPIA database, FC(log_2_)= −1.407. Significantly down-regulated signals in both the studies were FLNA, KLF4, FGFR2, IL6R, TSPAN7, WNT5B, HDAC4, CDKN2B, and COL4A5.

**Figure 4:**
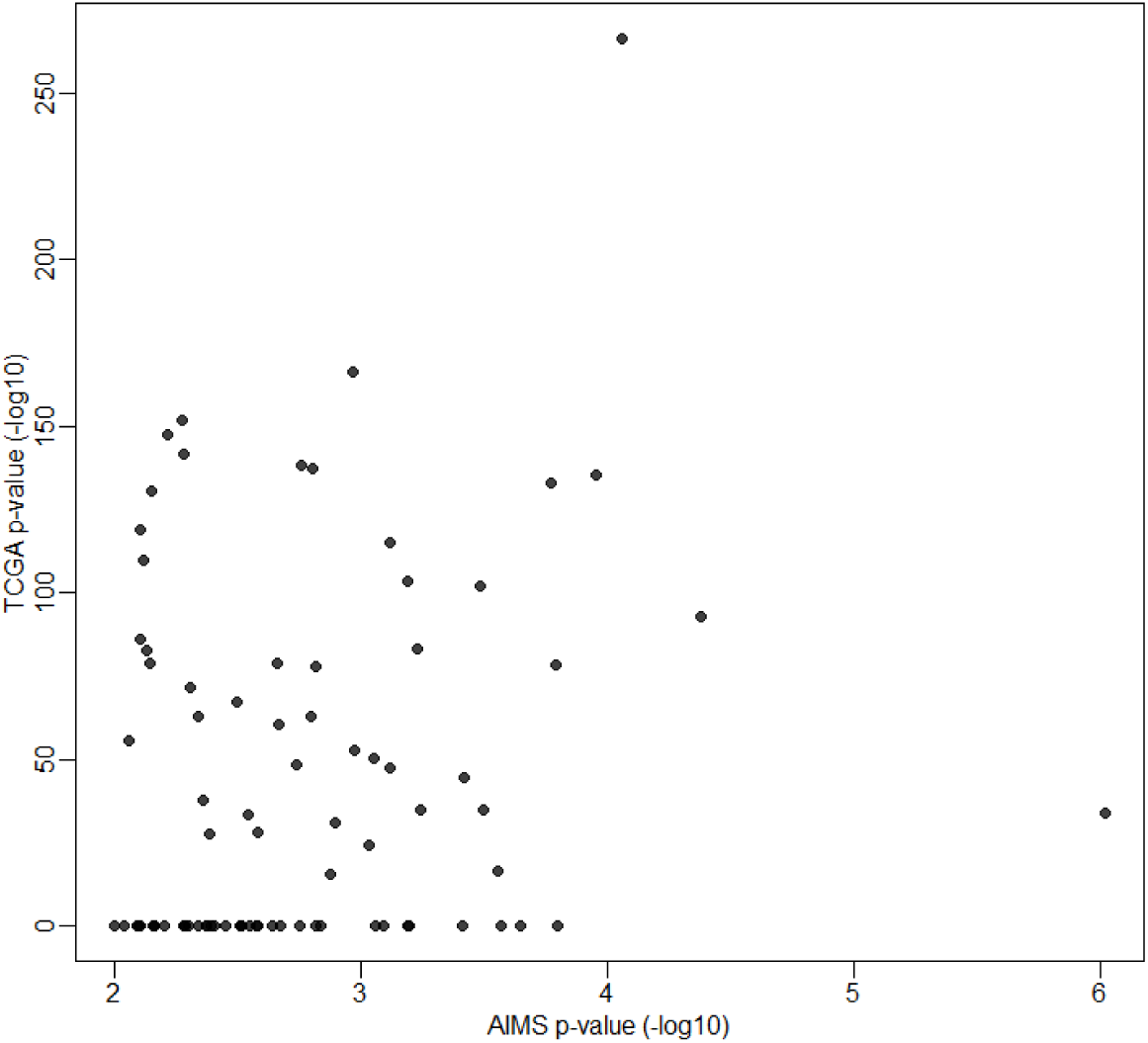
Comparison of 83 significant genes in the current study correlated with TCGA COAD expression data analyzed by GEPIA. The X-axis shows the *p*-values (-log_10_) of the current study and the Y-axis depicts the *p*-values (-log_10_) from the TCGA COAD data. 45 out of 83 genes had a significant difference in TCGA, while 38 genes were not significant in the TCGA data (as zero values in the graph).

#### TIMER

The list of highly significant 83 genes obtained from the Nanostring was fed into the TIMER TCGA COAD correlation module. Correlation module showed expression scatter plots between a pair of genes from the list in TCGA COAD cancer type with Spearman’s rho value with statistical significance (Figure 5a). Partial correlation with age and tumor purity was also analyzed in the given set of gene samples. Correlation data is given in Supplementary Dataset file S4. As shown in the correlation matrix, Figure 5a,from TIMER and the chord diagram, Figure 5b,expressions of COL1A1, COL11A1, COMP, INHBA, and SFRP4 had a score of 4.5,and SPP1 had a score of 3.5, all of them were found to be significantly correlated (Figure 5c). While DDIT4, RNF43, and ETV4 had a very low score (0.0 – 0.5) suggesting a low correlation. From the correlation heatmap (figure 5c), ETV4 and RNF43 expressions were significantly correlated to each other according to cluster analysis of the TIMER correlation database.

**Figure 5.**
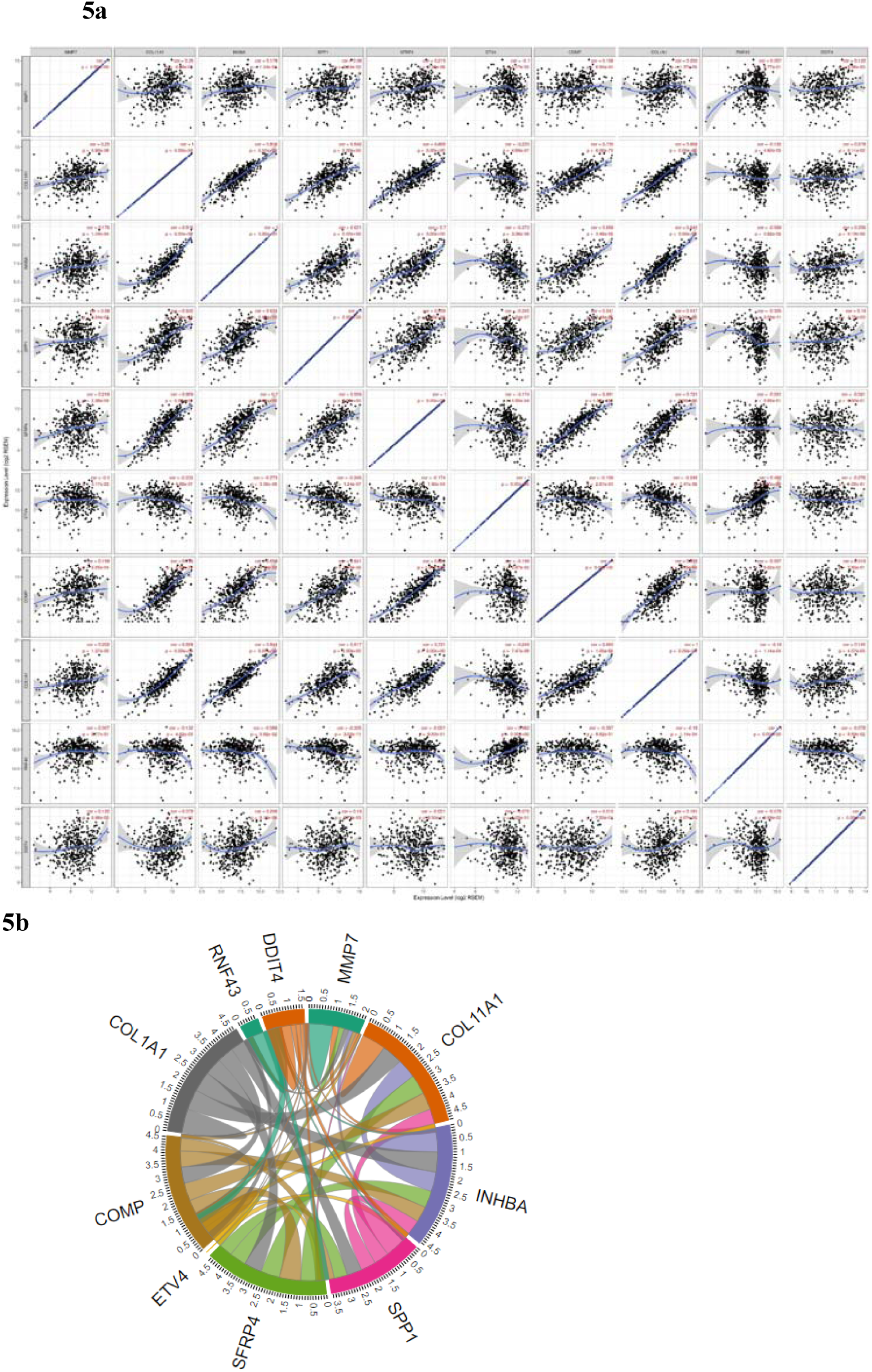
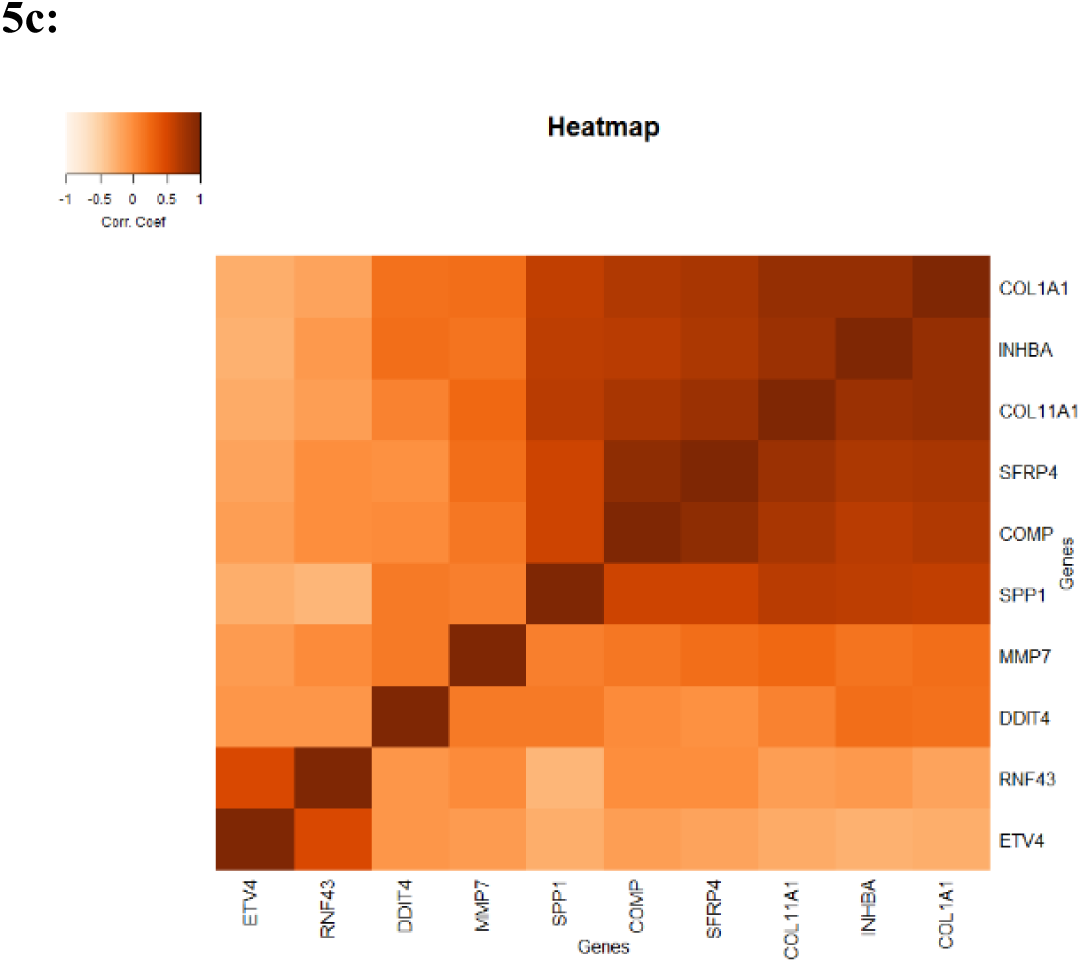
a: Co-expression correlation scatterplot of top 10 upregulated genes in the study having log_2_(FC) > 1 from the TCGA TIMER database. The correlation coefficient and the *p*-value are given in the top right corner of each of the scatterplot. As seen from the image, highly correlated signals would have a higher correlation. **5b: Chord diagram showing the correlation of TIMER co-expression data**. Top 10 genes that are correlated in the TIMER TCGA COAD database are arranged according to their correlation. The overall contribution of the correlation coefficient was found to be highest among six genes (COL11A1, COL1A1, COMP, SPP1, SFRP4, and INHBA; a score of 3.5 – 4.5). Genes that contributed less to overall correlation were DDIT4, RNF43, and ETV4 (score of 0.0-1.5). **5c: Heatmap showing the co-expression of the top 10 genes in the TIMER database**. Added to the high co-expression correlation with the six genes (COL11A1, COL1A1, SPP1, SFRP4, COMP, and INHBA), there is a separate correlation between RNF43 and ETV4 in a different cluster. Details are described in the main body of the text.

### Network Analysis

The top 20 highly significant genes from the Nanostring expression analysis were fed into the STRING.db online tool to explore the possible protein-protein interactions among these gene signals. The ranking of each of these proteins was determined from the fold-change values obtained from the differential expression ratio data from the current study. As illustrated in figure 6,eleven genes from the top 20 were found to be interacting with each other according to several databases included in STRING.db (gene co-occurrence and gene-neighborhood). Detailed list of interactions is given in Supplementary Dataset File S6). These were INHBA, COL1A1, COL11A1, COMP, SFRP4, SPP1, MMP7, and WNT2. These were indirectly connected to KLF4, CDKN2B, and ETV4. Among these, experimentally proved interactions were projected in three clusters. These were 1) INHBA, COL1A1, COL11A1, and COMP, 2) SFRP4, WNT2, SPP1, MMP7, KLF4, and CDKN2B, and 3) CACNA1H and CACNA2DH. Protein homology was found between COL1A1 and COL11A1, SFRP4 and WNT2, MMP7 and SPP1, and FLNA and FLNC. Interactions between different subunits of the interacting proteins (FLNC and FLNA, CACNA1H, and CACNA2D3) were also identified with the network analysis. There were no previous records of any gene fusions between them. Apart from these interactions, all the other genes from the top 20 genes were not shown to have any interaction with each other. The correlation cluster is seen with the TIMER TCGA database (and the genes clustered in the STRING.db network analysis was corresponding to many genes to each other.

**Figure 6:**
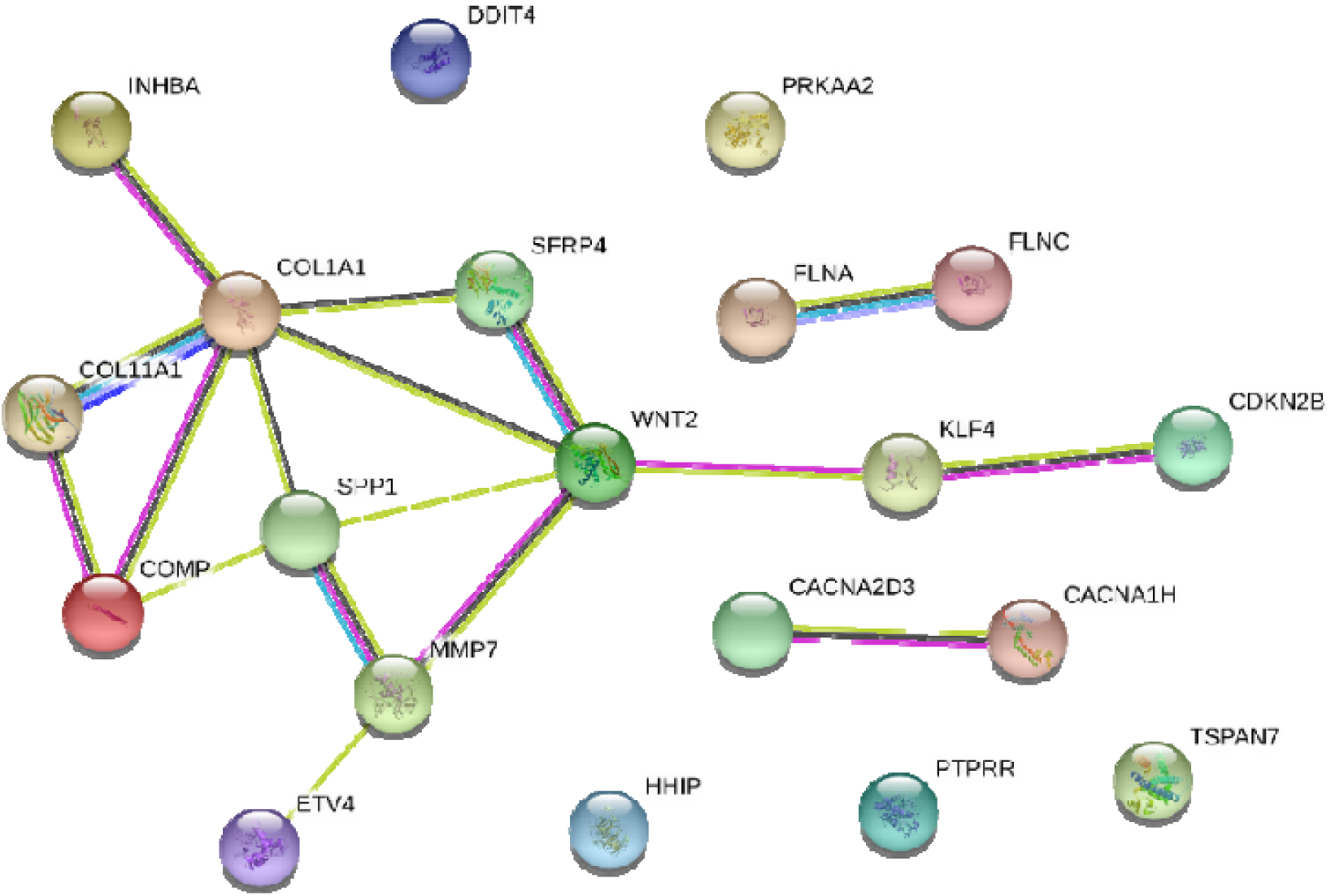
Network by STRING.db of the top 20 genes that were identified from the expression analysis. All the interactions were highly significant (FDR adj. *p*-value < 0.01). All colored nodes are the first shell of the interaction of each of the proteins. Edges represent protein-protein interaction, not necessarily connected. Color codes of interaction lines represent gene neighborhood (green), gene fusions (red), gene co-occurrence (dark blue), from curated databases (teal), experimentally determined (pink), text mining (yellow), co-expression (black), protein homology (light blue). The structures seen inside the nodes are the 3D structures of the known proteins.

### Association of immune cell infiltration with gene expression

Figure 7 shows a hierarchical clustering analysis of the correlation among the top 10 genes, identified by the Nanostring analysis from the current study cohort, with the immune cell infiltration data available in the TCGA COAD data obtained via TIMER. The immune cell correlation to the 83 significant genes are given in Supplementary Dataset File S5. Macrophages were aligned with the maximum number of genes within the given set. Five of the six highly co-expressed genes were associated with the tissue macrophage infiltration. Among them, COL11A1, COL1A1, INHBA, SPP1, SFRP4, and COMP were clustered with macrophages. Macrophages, CD4^+^T lymphocytes, dendritic cells as well as neutrophils had a typically similar association with a small variance across the different genes. However, the first three genes were clustered with CD4^+^T-cells, dendritic cells, as well as neutrophils. These signals were less associated with CD8^+^T cells as well as B-cells. This shows that the high expressions of the first six signals may have originated from the tumor infiltrates. Signals like MMP7 and DDIT4 were not associated with any specific cell type. However, ETV4 and RNF43 were clustered with tumor purity and away from all other infiltrating cell types. This suggests that the expressions of ETV4 and RNF43 might be represented by the tumor cells.

**Figure 7:**
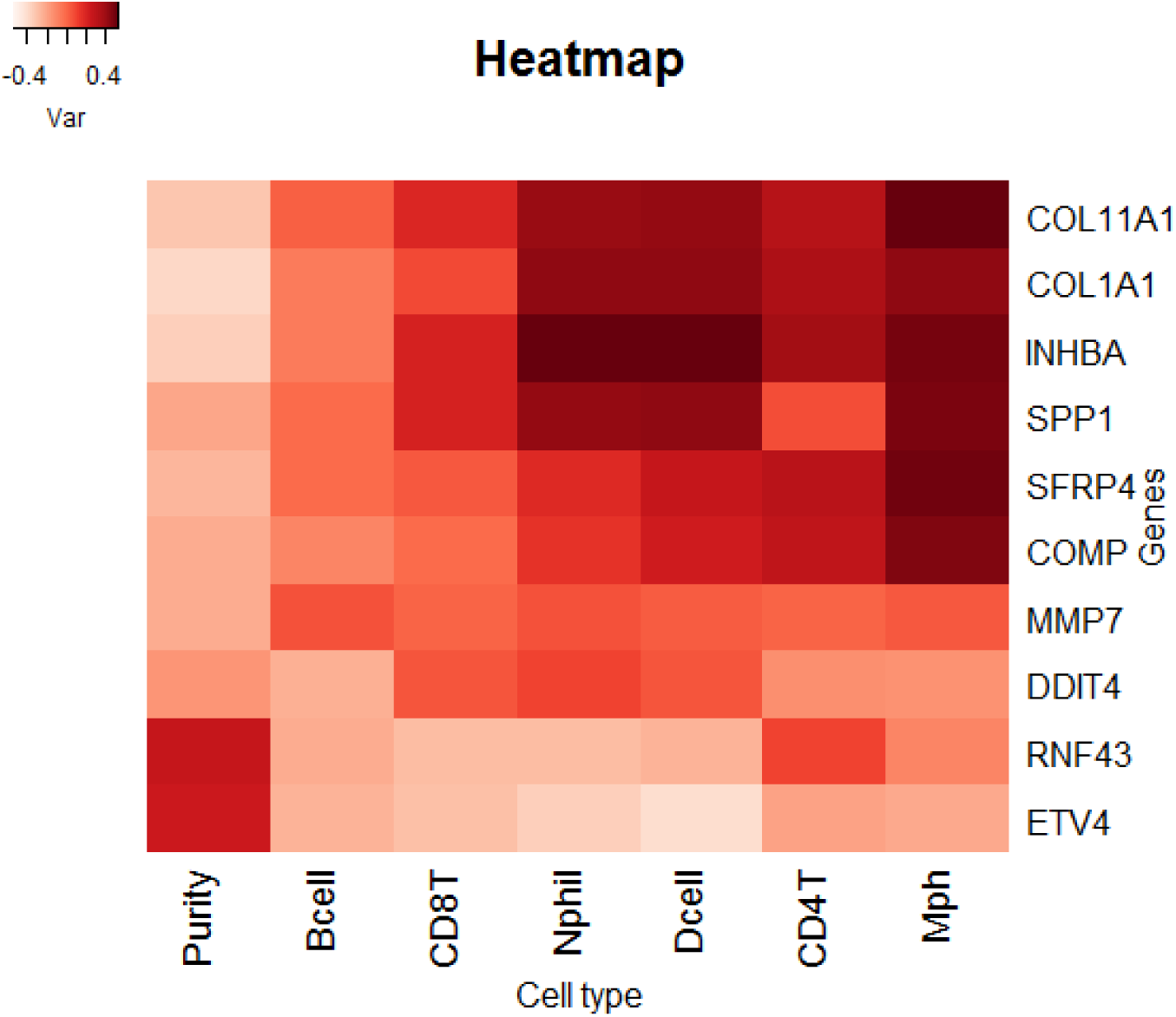
Comparison of the top 10 genes from the current study to immune cell infiltration in the TCGA TIMER database. Tissue Macrophages (Mph) are highly associated with the majority of the genes upregulated, followed by CD4-T lymphocytes, Dendritic cells (Dcell), Neutrophils (Nphil), CD8-T lymphocytes, and B-cells in the descending order. The tumor content (represented by “Purity”) is highly correlated with ETV4 and RNF43.

## Discussion

One of the major considerations of the large scale TCGA/Pan-Cancer Atlas studies is to correlate molecular pathogenesis data with the gross pathological and clinical parameters of cancerogenesis in different populations. The molecular classifications prescribed by the TCGA/ Pan-Cancer Atlas are derived from tumor samples of a wide variety of tissues at various levels of tumor stages, infiltrating cellular types, and different ethnic groups. Correlation of different data in TCGA to clinical medicine requires clinical molecular validation studies and analysis for their translational capacity in different populations. To that end, molecular signals from tumor samples of different populations that are annotated and stratified can be overlaid on to the TCGA at multiple data layers to examine the prognostic and pathogenic role of individual signals in a particular population. Using this method, the strength of the exhaustive analysis of TCGA can be applied directly to clinical implementation. In the current study, we have shown the first expression profile of colon adenocarcinoma in the South Indian population, a population that has not been widely represented in most of the previous large cohort studies. To validate these results, we showed preliminary methods of overlapping the results from the current study with TCGA data using three different methods. First is the comparison of the significant genes identified in the two datasets, second is the co-expression correlation between the genes in both datasets and thirdly is the comparison of differentially expressed genes in the study cohort with immune cell co-localization.

Among all the differentially expressed genes, in multiple group comparisons and the TCGA database by GEPIA and TIMER analysis, ETV4 was the most significant gene identified in our study. ETV4 expression was particularly high in tissues that have proficient MMR. This may be the first study to show a high correlation of ETV4 with MSS CRCs. ETV4 or ETS Variant Transcription factor 4 (E1A enhancer-binding protein – E1AF or Polyoma Enhancer Activator 3 Homolog – PEA3) is a known transcription factor, upregulated and activated in colon adenocarcinoma ^20-22^. Overexpression of ETV4 was shown in breast cancer via induction of HER2/Neu ^23,24^. ETV4 or other PEA3 factors (ETV1 and ETV5) were shown to initiate tumorigenesis by chromosomal translocation with other genes regions such as EWS in Ewing Sarcoma ^25^or with TMPRSS2 in prostate cancer ^26^. ETV4 has also been shown to upregulate the expression of MMP7 in many gastrointestinal tumors ^27^. Although in the current study, MMP7 expression was not highly significant across the samples, it showed the highest fold-change in DE (Figures 1). The reduced expression of ETV4 in MSI tumors could be due to the presence of dinucleotide GT repeats in the third intron of the *ETV4*gene, causing frame-shift mutation in the coding region. In MSS tumors, ETV4, along with the β-catenin-induced Tcf-mediated *MMP7*promoter activation in intestinal and colon cancer cells ^28^. However, a specific driving factor for the upregulation of ETV4 in MSS tumors is unclear from previous studies. ETV4 binding motif is located in the upstream-144 of the *MMP7*promoter region. However, the domain of the ETV4 that binds to the promoter region is unknown yet. Matrix-metalloproteinase-7 (MMP7) is a commonly upregulated gene in colon cancers ^27,29-31^. ETV4 induced MMPs (MMP1 and MMP7) are involved in collagen catabolism, both being molecular markers for the epithelial-mesenchymal transition ^32^. In the current study, though ETV4 was found to be elevated in MSS samples, it was also elevated in cases of MSI with TILs. As an offshoot of ETV4 upregulation mediated MMP-7 activation, proMMP-9 can be activated to attach to a cellular G-protein coupled receptor mechanism for the activation of the Neuraminidase-1 enzyme, which causes cleavage of α2,3-sialic acid causing immune recognition has been recently identified as a mechanism in many cancers^33,34^. Though post-translational modification and activation of ETS proteins by either phosphorylation ^35^or transcriptional co-activation via acetylation or sumoylation ^26^have already been demonstrated, signals that induce the expression of ETV4 in MSS tumor cells are still not completely understood.

The Nanostring PanCancer panel includes many targets that are associated with tumor microenvironment and invasion. The current study has depicted a cluster of genes that are upregulated in the tumors, compared to normal tissues, although not clustered along with the tumor cells. Signals included are COL1A1, COL11A1, SPP1, COMP, SFRP4, and INHBA. Numerous studies have shown the proliferative role of collagen in gastric ^32,36^, breast ^37^, and colon cancers ^38,39^. Previous studies have also demonstrated that the upregulation of MMPs (MMP7 and 9) along with other collagen proteins are important for tumor vascular invasion ^40^. Along with MMPs and other signaling markers of Epithelial-Mesenchymal Transition (EMT), COMP ^41^and SFRP4 ^42^were also found to be upregulated in colon cancers and associated with poor overall survival. Inhibin β A (INHBA), is a member of Transforming Growth Factor β (TGF-β) superfamily, whose increased expression has been shown earlier in colon adenocarcinoma cells ^43^, and was recently demonstrated to have a prognostic significance as well ^43^.

The clustering of genes from the network analysis (Figure 5) and TIMER correlation analysis (Figure 6), in the current study, showed two distinct clusters of gene signals, possibly residing in two different cell types/groups of the tumor. The first cluster contained COL1A1, COL11A1, COMP, SFRP4, SPP1, and INHBA which were mainly associated with samples with higher infiltration of tumor-associated macrophages (TAM), TILs (CD4^+^cells), dendritic cells and neutrophils in TGCA TIMER data. The second cluster contained ETV4 and RNF43 whose expression was associated with the purity of tumor cells in TIMER data. MMP7 and DDI4 were mostly associated with cells that were not associated with high purity of tumor cells. This may implicate that the ETV4 and RNF43 might be directly expressed by tumor cells, while others were expressed in infiltrating non-tumor cells. Since the proliferation of both tumor cells and infiltrating immune cells are required for the tumor evolution and progression, the proliferative role of these signals may apply to these cell types. Though the cell types associated with these signals were previously reported in several studies ^44,45^, the signals correlated in the current study have not been associated with any of the tumor immune cells. Hence, these results may imply that the current set of signals, might present in the immune cells as well. The high-throughput expression analysis approach is limited in its application, in this context, to delineate the location of these signals in a complex tumor microenvironment. More elaborate studies such as immunohistochemical colocalization experiments are required to identify the cellular location of these signals in tumors.

## Conclusions

We have shown, in this study, a method to bridge the gap between the large cohort molecular pathogenesis discovery studies, such as TCGA, to low-throughput and uncharacterized pathological tumor samples from routine clinical medicine, often from unique or highly focused ethnic population. We explored a high throughput expression analysis using the Nanostring PanCancer pathway panel in 11 stage-II colon and rectal adenocarcinoma tumor from South Indian (Kerala) population. 83significant genes enriched cell cycle-apoptosis, chromatin modification, and DNA damage repair pathways. The signals revealed in the study were overlapped with TCGA at three different data layers. Among the top 20 differentially expressed genes with largest FC, two sets of gene signals were clustered, according to the comparative co-expression analysis using GEPIA and protein interaction network analysis. Similar segregation of signals was found in tumor immune infiltration status according to TIMER. The former mRNA signal cluster was found to be associated with macrophages, CD4^+^T cells, dendritic cells, and neutrophils. Of the two signals in the second cluster, one signal, ETV4, was significantly higher in MSS tumors. The significance of the high expression of ETV4 with respect to MSS and its associated cause and/or effect in Indian CRC is an unexplored area to date. The signals revealed in this study need to be further validated in larger patients’ samples in the same population in order to check pathogenic role and predictive and prognostic utility. We think that methods like this, if implemented and validated in different populations in larger numbers, would help in the identification of more precise prognostic predictive markers and effective agents that can counter the pathogenesis to provide better patient care and effective intervention.

## Methods

### Samples

Archived tumor tissues, which were formalin-fixed paraffin-embedded (FFPE) during the previous year, were utilized for the study. We selected 11 tumor tissues of Stage II CRC, and four normal tissues for the Nanostring nCounter analysis. Institutional scientific and ethics committee approval was obtained before the study. All the samples were previously diagnosed by histopathology as well as immunohistochemistry and/or MSI-PCR for diagnosing MMR deficiency. The normal tissues were obtained from the normal regions of the tumor tissue, which were also histopathologically identified. The study was conducted as per the guidelines of the Institutional Ethics Committee of Amrita Institute of Medical Sciences (AIMS). The project was presented to the Institutional Regulatory Board (IRB), which comprises the Scientific Review Committee and the Institutional Ethics Committee. All the tumor samples included in this study were deidentified. The tissues were obtained as part of the usual diagnostic and treatment policies of the institution. This study did not change the clinical management of those patients. Considering all these facts, the committee decided that there is no need for separate consent from each patient. The study was reviewed and approved by the IRB on Nov 24^th,^2017 (IRB-AIMS-2017-124).

### Sectioning and Microdissection

Serial 5μm histological sections of FFPE tissue block of normal and tumor were prepared. The first and fifth sections were stained with hematoxylin and eosin (H&E), sandwiched tissue levels were mounted unstained on slides, verified by light microscopy for each block using levels 1 & 5, and areas to be micro-dissected were marked. Normal or tumor tissue was micro-dissected from the unstained slides for each case by overlaying the unstained slides in a laminar flow tissue culture hood. Alternatively, one section from the trimmed block is sampled for a tumor and a normal area that is marked in the block. The block is cut in those areas and re-embedded in separate blocks and sectioned into separate sample tubes for further processing.

### Nanostring nCounter assay

The NanoString nCounter^®^Analysis System delivers direct, multiplexed measurement of gene expression, providing digital readouts of the relative abundance of hundreds of mRNA transcripts simultaneously ^46,47^. The nCounter Analysis System is based on gene-specific probe pairs that are hybridized to the sample in solution. The protocol eliminates any enzymatic reactions that might introduce bias in the results. The Reporter Probe carries the fluorescent signal; the Capture Probe allows the complex to be immobilized for data collection. Up to 550 pairs of probes, specific for a set of genes are combined with a series of internal controls to form a CodeSet.

The assay was conducted by a contract research organization (Theracues Pvt Ltd, Bangalore, India) on a pay-by-service basis. According to the protocol from the organization, briefly, three-to-four 5μm FFPE samples sections were taken for RNA extraction using commercial FFPE nucleic acid isolation kit (Roche Molecular Diagnostics). RNA was quantified using Nanodrop and analyzed for fragment distribution using a bioanalyzer (Agilent). Considering the profile on the bioanalyzer ∼140 ng of RNA was used for each probing assay. Hybridization with PanCancer Pathway Panel codeset was performed for 18 hrs. Post-hybridization, this was separated on nCounter sprint machine on a microfluidic cartridge. The data were normalized, and fold change was generated using nSolver 4.0 software.

### Analysis

For Nanostring expression assays, data from NCBI Gene Expression Omnibus profiles (http://www.ncbi.nlm.nih.gov/geo/) from independent cancer gene sets (accession numbers such as GSE14333, GSE35602, etc.) were used for expression analysis. To find the fold-change, a ratio of median values of the probeset expression in each of the tumor samples to the median value of the probeset expression in the normal tissue was identified. The median values of the probe set expression, after standardization with standard expression sets, and fold-change values were drawn in data matrices, and two-dimensional hierarchical clustering analysis with different Euclidean distance clustering methods was employed using appropriate package modules in R statistical program ^9^. Advanced Analysis includes the *Pathway score*and *Differential Expression*Analysis. Nearly 770 genes were probed in a single PanCancer Pathway hybridization chip, including about 606 genes were from thirteen canonical pathways, and about 125 genes were cancer driver genes. Approximately 50 genes were included as reference and housekeeping genes. The fluorescence count obtained from the nCounter machine (cutoff value of ≥ 40) was normalized according to the positive controls and internal housekeeping reference genes. Among the four tumor-normal paired samples, included in the pilot study, the Differentially expressed (DE) genes in the tumor were calculated by taking the geometric mean of the four normal tissue as the denominator and the geometric mean of all the tumor samples as the numerator. The fold changes were calculated from this ratio data. If ratio > 1; then Foldchange = Ratio, if ratio < 1; then Foldchange = −1*(1/Ratio). The fold changes were then reported. The Fold Change data were filtered for Fold changes ≥ 1.5 & ≤ −1.5. The genes with counts ≥ 40 in normal or tumor samples were retained. The final list of genes was then marked in red for downregulation and green for upregulation. The top 10 up- and down-regulated genes were used to generate the principal component analysis (PCA) plot. Genes were tested for differential expression in response to each selected sample. For each gene, single linear regression was fit using all selected covariates to predict expression to eliminate confounding results due to measured covariates, and to isolate the independent association of each covariate with gene expression, measuring each variable’s association with a gene after holding all other variables constant. Evaluation of differential expression values by group difference between microsatellite instability status, infiltration status as well as the anatomical location were analyzed using Student t-test analysis assuming unequal variance. The Pathway analysis was done for each sample separately using DE genes (DE Call − Yes) using WEB-based GEne SeT AnaLysis Toolkit (WebGestalt)^48^. The Enrichment was carried out for functional databases Kyoto Encyclopaedia of Genes and Genomes (KEGG) ^49^and Panther Pathways ^50^using the Gene Ontology summary database (GOSlim). From the set of genes obtained from the WebGestalt tool, further statistical analysis using PCA was performed for dimension reduction. PCA was done using ClustVis ^51^, an R-program based component factor analysis tool.

### Network Analysis

STRING database (string-db.org) is an integrated, publicly available resource for identifying integrated protein-protein interaction networks in biology, developed by STRING consortium ^52^. The interaction network was built on laboratory experiments in protein-protein interactions, conserved co-expression data, genomic context predictions, and text mining from previous research. The genes that were found to be highly significant across the samples and maximal log2 fold-change (either upregulated or downregulated) from the Nanostring nSolver analysis were selected by the appropriate R-program module. These genes, with their log2 fold-change values, were uploaded into the STRING-db.v.11 online search tool, in the “Proteins with Values/Ranks” tab.

### Co-expression analysis

The network identified from the STRING database was further evaluated by co-expression analysis by GEPIA and TIMER databases. GEPIA is an online tool (http://gepia2.cancer-pku.cn/) to explore RNA sequencing expression analysis of 9736 tumors and 8587 normal samples from the TCGA and GTEx projects using standard processing pipeline ^53^. One-to-one gene expression analysis is possible with GEPIA. Using both databases, the correlation of each of the selected genes, found to be expressed together from the test samples, were analyzed for their co-expression correlation coefficient in the TCGA COAD using TIMER database (https://cistrome.shinyapps.io/timer/). The spearman correlation coefficient obtained from the TIMER is compared against the network analysis results obtained from STRING.db.

### Immune cell infiltrate Analysis

TIMER (cistrome.shinyapps.io/timer) is a web-based database and computational tool to statistically analyze tumor-infiltrating immune cells from gene expression profiles ^19^. The tool compares the immune cell infiltrates such as B-cells, CD4^+^T cells, CD8^+^T cells, neutrophils, macrophages, and dendritic cells from the TCGA data, which contain more than 10,000 samples from 32 different cancer types. The selected genes from the study samples were compared against the TIMER immune cell infiltrate to identify the type of immune cells that were maximally expressed in the TCGA COAD cohort from the given set of unique genes identified from the local population. The correlation coefficient of each cell type in the TIMER database against the top enriched genes from the current study is analyzed and clustered using hierarchical clustering analysis to identify the set of genes that may have an association with a particular cell type. Clustering analysis was done using base functions such as *hclust*, and *dist*functions in the R program.

## Data availability

Supplementary and other data are publicly available at the Open Science Forum website. The data can be accessed at the weblink: https://osf.io/tcdk9/.

## Acknowledgments

The authors would like to thank the senior and junior faculty staffs, office and laboratory staffs of Departments of Medical Oncology, Pathology, Molecular Biology, Biochemistry, Health Sciences Research and Molecular Oncology Laboratory of Amrita Institute of Medical Sciences, Kochi for the coordination of resources for the conduct of research. The authors would also like to thank the staff and CEO Mr. Gopalakrishna Ramaswamy of Theracues Innovations Pvt Ltd, Bangalore, India for the coordination and conduct of Nanostring assays. The study was supported by an internal seed grant by Amrita Vishwa Vidyapeetham to PSA.

## Author contributions

PSA has designed the experiments with RAJ, VM, RRP, and KP. VM and DMV reviewed the project and further modified. Conduct of the research was done by PSA and RAJ. Data analysis was done by PSA. The manuscript was written by PSA and RAJ, reviewed by VM, RRP, KP, and DMV.

